## Supplementary data for "Triggering MSR1 promotes JNK-mediated inflammation in IL-4 activated macrophages"

**Table EV3**. Ubiquitylated proteins and their intensities identified in Halo-TAB2 TUBE pull-downs from phagosomes of M2(IL-4) macrophages

| Gene name | Protein | Intensity TAB2-mut | Intensity TAB2 | Ubiquitylated position |
|---|---|---|---|---|
| Ubc | Polyubiquitin-C | $2.7\times10^{4}$ | $3.9\times10^{5}$ | 63;11;48;1;29;6;27;33 |
| **Msr1** | **Macrophage scavenger receptor 1** | $9.7\times10^{4}$ | $1.3\times10^{7}$ | **27** |
| Slc43a2 | Large neutral amino acids transporter small subunit 4 | $2.2\times10^{6}$ | $1.2\times10^{7}$ | 283;293;402;557 |
| Ifitm3 | Interferon-induced transmembrane protein 3 | $6.8\times10^{4}$ | $3.9\times10^{6}$ | 24 |
| Rnf13 | E3 ubiquitin-protein ligase RNF13 | $1.3\times10^{5}$ | $7.7\times10^{6}$ | 233 |
| Csf1r | Macrophage colony-stimulating factor 1 receptor | $3.3\times10^{6}$ | $8.0\times10^{7}$ | 572;810;625;584 |
| C5ar1 | Complement component 5a receptor 1 | $5.0\times10^{6}$ | $1.8\times10^{7}$ | 334 |
| Slc38a2 | Sodium-coupled neutral amino acid transporter 2 | $3.5\times10^{5}$ | $5.0\times10^{6}$ | 38;33 |
| Serinc3 | Serine incorporator 3 | $8.6\times10^{4}$ | $5.0\times10^{5}$ | 363 |
| Slc7a2 | Isoform 2 of Low affinity cationic amino acid transporter 2 | 0 | $3.5\times10^{6}$ | 449;632;654 |
| Tyrobp | TYRO protein tyrosine kinase-binding protein | 0 | $5.4\times10^{5}$ | 82 |
| Lyn | Tyrosine-protein kinase Lyn | $8.6\times10^{6}$ | $2.0\times10^{7}$ | 20 |
| Unc93b1 | Protein unc-93 homolog B1 | $4.1\times10^{6}$ | $1.4\times10^{7}$ | 197;582 |
| Itm2b | Integral membrane protein 2B | $1.6\times10^{6}$ | $1.1\times10^{7}$ | 13 |
| Slc6a6 | Sodium- and chloride-dependent taurine transporter | $6.7\times10^{5}$ | $3.1\times10^{7}$ | 611;15;37 |
| Lat2 | Linker for activation of T-cells family member 2 | 0 | $1.0\times10^{6}$ | 39;84 |
| Tfrc | Transferrin receptor protein 1 | $7.8\times10^{6}$ | $2.2\times10^{7}$ | 53;39 |
| Abcg1 | ATP-binding cassette sub-family G member 1 | $1.3\times10^{6}$ | $9.9\times10^{6}$ | 55 |
| Slc40a1 | Solute carrier family 40 member 1 | $1.9\times10^{4}$ | $2.2\times10^{5}$ | 269 |
| Itch | E3 ubiquitin-protein ligase Itchy | $1.9\times10^{5}$ | $8.1\times10^{6}$ | 192 |
| Piezo1 | Piezo-type mechanosensitive ion channel component 1 | $3.9\times10^{4}$ | $5.3\times10^{6}$ | 1914 |
| Ednrb | Endothelin B receptor | $3.4\times10^{6}$ | $9.5\times10^{6}$ | 417 |
| Vamp8 | Vesicle-associated membrane protein 8 | $1.0\times10^{5}$ | $2.7\times10^{6}$ | 47 |
| Mgl2 | Macrophage galactose N-acetyl-galactosamine | $1.0\times10^{7}$ | $8.7\times10^{7}$ | 16 |
| Hspa8 | Heat shock 70 kDa protein 8 | $1.3\times10^{7}$ | $6.1\times10^{7}$ | 524 |
| Slc23a2 | Solute carrier family 23 member 2 | $7.9\times10^{4}$ | $3.2\times10^{6}$ | 11 |
| Evi2a | Protein EVI2A | 0 | $1.1\times10^{6}$ | 192;221 |
| Itfg3 | Protein ITFG3 | $5.8\times10^{4}$ | $1.8\times10^{6}$ | 36;38 |
| Emr1 | EGF-like module-containing mucin-like hormone receptor-like 1 | $3.6\times10^{6}$ | $1.9\times10^{7}$ | 743 |
| Slc12a4 | Solute carrier family 12 member 4 | $8.4\times10^{4}$ | $4.3\times10^{6}$ | 988 |
| Slc3a2 | 4F2 cell-surface antigen heavy chain | $3.5\times10^{6}$ | $3.2\times10^{7}$ | 42 |
| Ttyh3 | Protein tweety homolog 3 | 0 | $1.3\times10^{7}$ | 508 |
| Clec4a3 | C-type lectin domain family 4 member a3 | 0 | $1.6\times10^{5}$ | 13 |
| Colec12 | Collectin-12 | 0 | $2.3\times10^{5}$ | 2;17 |
| Ms4a6d | Membrane-spanning 4-domains subfamily A member 6D | 0 | $2.4\times10^{5}$ | 174 |
| Slc5a3 | Sodium/myo-inositol cotransporter | $1.6\times10^{5}$ | $1.7\times10^{6}$ | 596 |
| Trpv2 | Transient receptor potential cation channel subfamily V member 2 | $3.5\times10^{6}$ | $2.3\times10^{7}$ | 51 |
| Slc43a3 | Solute carrier family 43 member 3 | 0 | $2.6\times10^{5}$ | 244 |

### Supplementary Figure Legends

Figure EV1

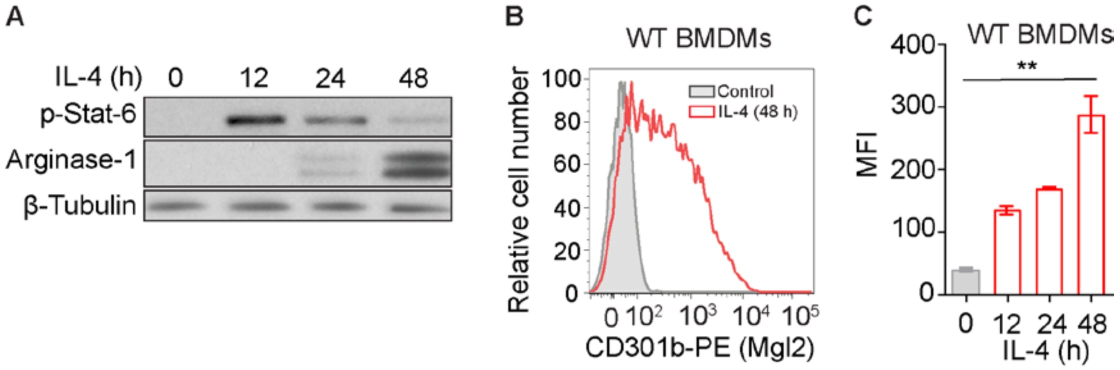


***Figure EV1: Validation of IL4-activation.***

(A) Immunoblot analysis (IB) of IL-4- induced M2 markers including pStat-6 and Arginase-1. Macrophages were fully activated after 48 h as shown by strong induction of Arginase-1. Tubulin serves as loading control. (B) The cell surface levels of CD301b/Mgl2 in response to IL-4 stimulation. (C) Quantification of mean fluorescence intensity (MFI) of cell surface levels of CD301b/Mgl2 in response to IL-4 stimulation.

Figure EV2

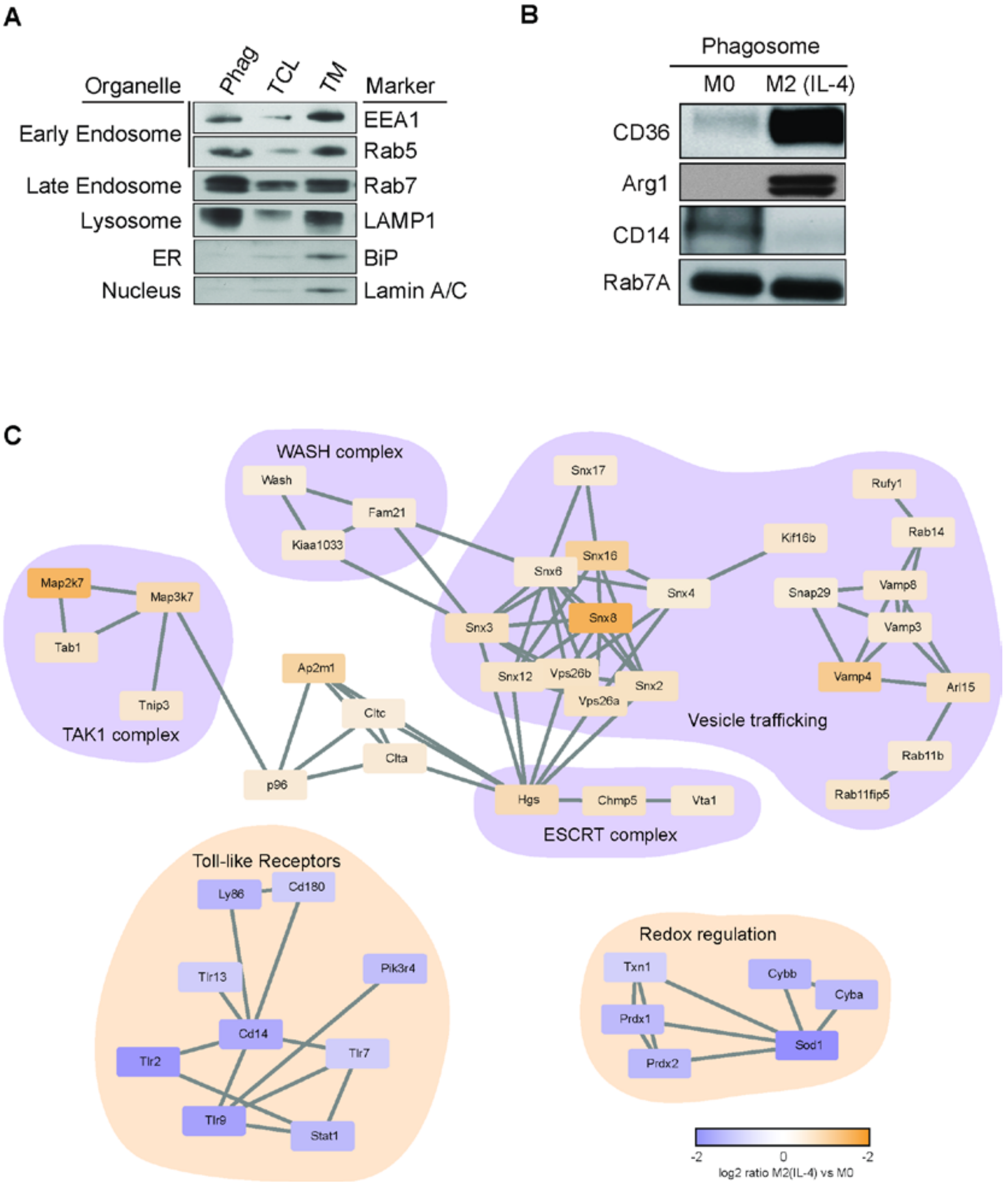


***Figure EV2: Validation of phagosome proteomics data and specific regulated phagosome protein complexes.***

(A) IB of markers of early and late endosomes (EEA1, Rab5a and enriched on phagosomes, while nuclear and ER markers (Lamin A/C and BiP) are more abundant in total cell lysates (TCL) and total membrane (TM), respectively. (B) IB analysis validating selected proteins changing in proteomics data. (C) Protein–protein interaction network of phagosome proteome using the STRING database v10 (Szklarczyk D, et al, Nucleic Acid Res, 2015). Subnetworks positively affected by phagocytosis in M2(IL-4) macrophages were vesicular trafficking and signalling, while proteins associated with innate immune response and ROS were down-regulated. Proteins involved in vesicular trafficking and signalling, Redox regulation and Toll like receptor complexes are displayed. (A-B) Representative blots of two biological replicates.

Figure EV3

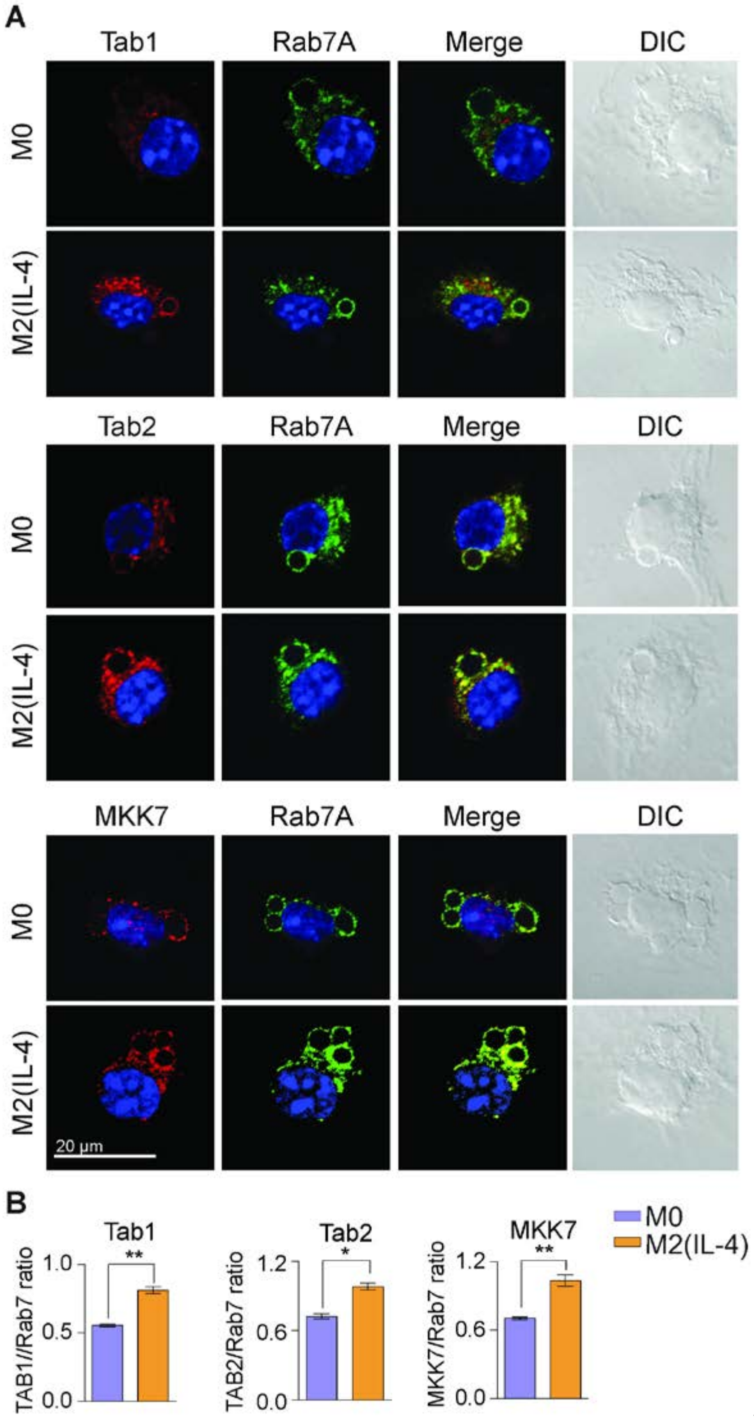


***Figure EV3: Immunofluorescence of TAB1, TAB2 and MKK7 on M0 and M2 phagosomes.***

(A) Immunofluorescence micrographs showing co-localisation of TAB1, TAB2 and MKK7 proteins with Rab7a, a phagosomal marker, around 30 min phagosomes in M0 and M2(IL-4) macrophages. (B) Corresponding quantification of co-localisation of TAB1, TAB2 and MKK7 with Rab7a in M0 and M2(IL-4) macrophage phagosomes, plotted as a ratio of the fluorescence of the target protein to Rab7a fluorescence localized to phagosomes. Scale bar is 20 µm. Data are shown as mean ± SEM from three independent experiments. $^{**}p < 0.01$, $^{***}p < 0.001$; (Student's T-test).

Figure EV4

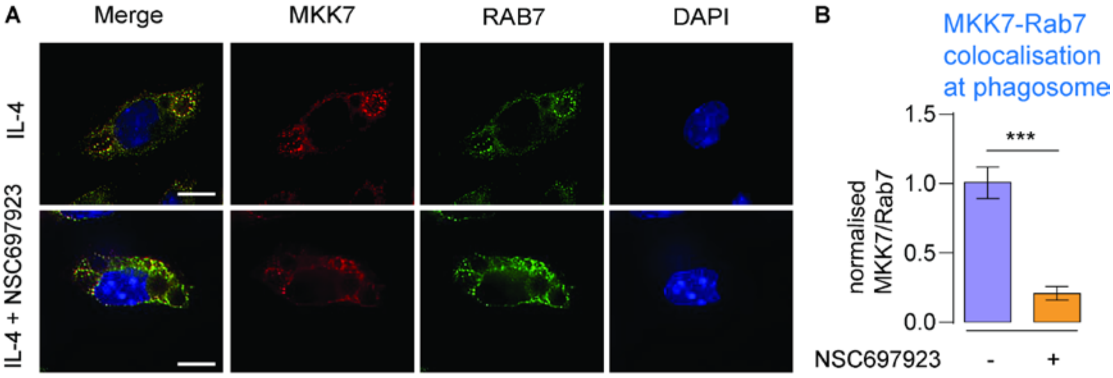


***Figure EV4: Immunofluorescence of TAB1, TAB2 and MKK7 on M0 and M2 phagosomes.***

(A) Immunofluorescence micrographs showing co-localisation of MKK7 protein with Rab7a, a phagosomal marker, around 30 min phagosomes in M2(IL-4) macrophages. Co-localisation is reduced upon treatment with the UBC13 inhibitor NSC697923. (B) Corresponding quantification of co-localisation of MKK7 with Rab7a in M2(IL-4) macrophage phagosomes ± treatment with NSC697923, plotted as a ratio of the fluorescence of MKK7 to Rab7a fluorescence localized to phagosomes. Scale bar is 10 µm. Data are shown as mean ± SEM from three independent experiments. $**p < 0.01$, $***p < 0.001$; (Student's T-test).

Figure EV5

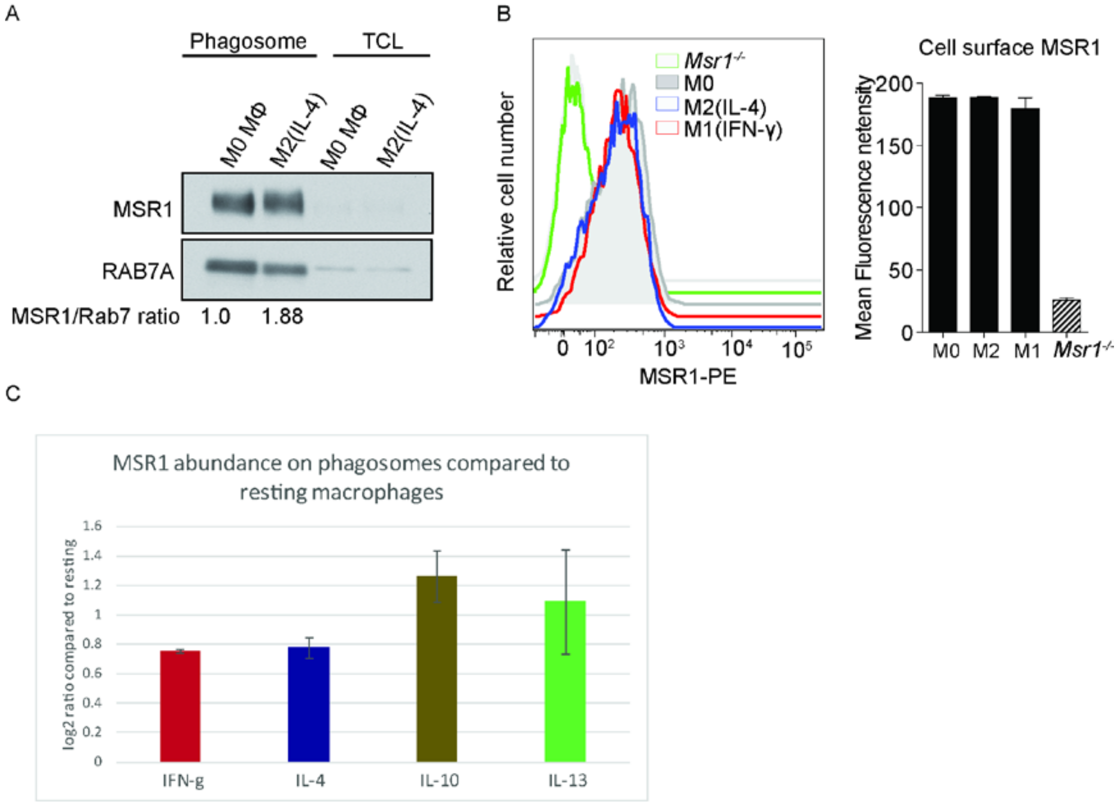


***Figure EV5: Identification of MSR1 expression and K63 polyubiquitylation on phagosomes in activated macrophages.***

(A) Immunoblot of MSR1 showing light enrichment of the non-ubiquitylated form of the protein on phagosomes in M2(IL-4) macrophages compared to M0 macrophages. (B) Activation does not affect MSR1 surface expression. Flow cytometry analysis and corresponding quantification shows equal cell surface expression of MSR1 in M0, M2(IL-4) and M1(IFN-γ) macrophages. Data are shown as mean ± SEM from three independent experiments. MFI (Median Fluorescence Intensity). (C) Proteomics protein abundance data for MSR1 on 30 min old phagosomes from IFN-γ (100 U/ml, 24 hrs), IL-4 (20 ng/ml; 48 hrs), IL-10 (10 ng/ml; 48 hrs) and IL-13 (20 ng/ml; 48 hrs) activated macrophages compared to phagosomes from untreated/resting macrophages. The data shows that MSR1 is induced on phagosomes upon all of these treatments. Data are shown as mean ±standard deviation from three independent biological experiments.

Figure EV6

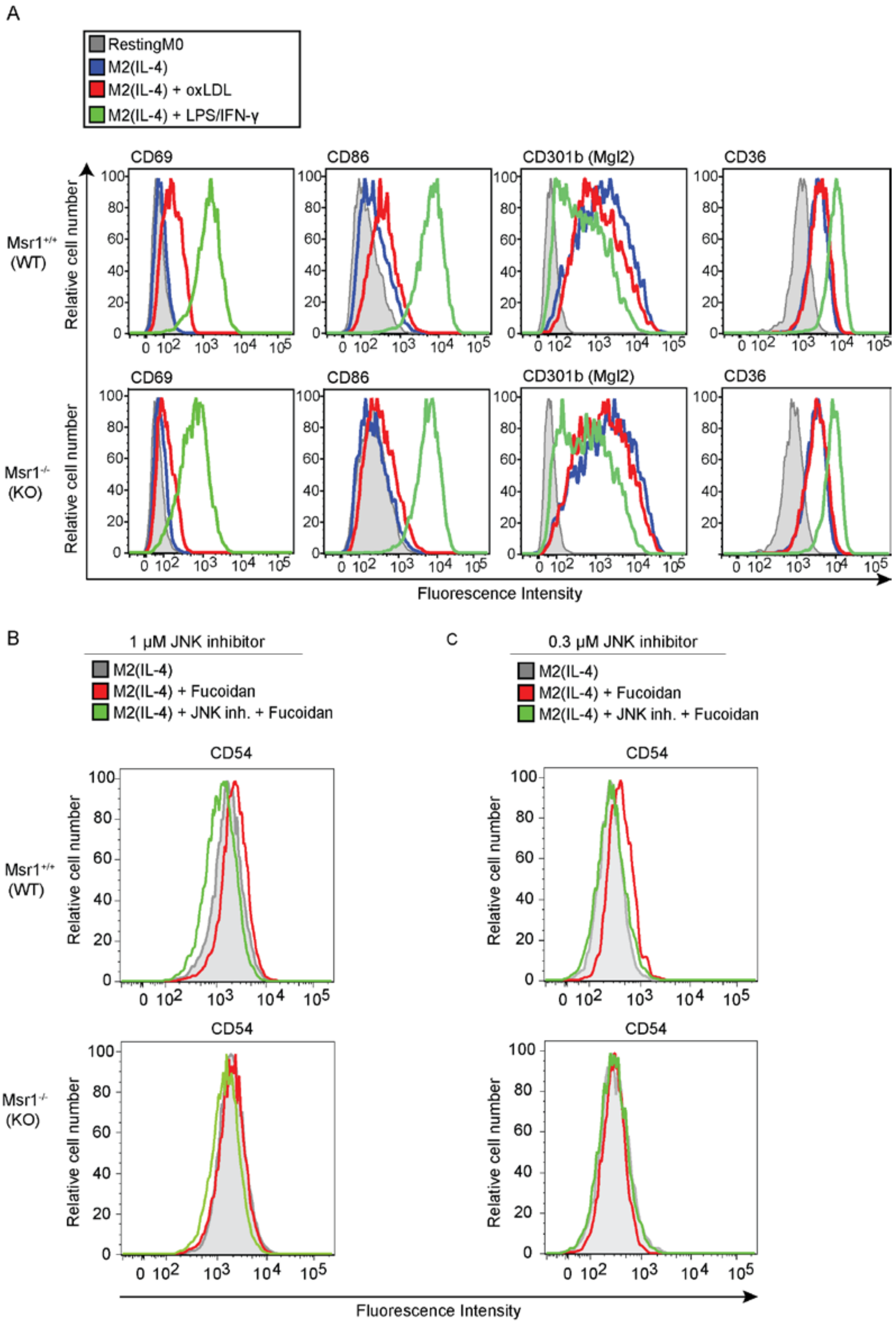


***Figure EV6: oxLDL induces a MSR1-dependent pro-inflammatory switch and JNK inhibition reverses MSR1-dependent pro-inflammatory activation.***

(A) Flow cytometry analysis of cell surface expression of CD69, CD86, CD301b and CD36 of M2(IL-4) followed or not by either oxLDL or LPS/IFN-γ treatment. (B-C) Inhibition of JNK by 1 and 0.3 μM (C) of JNK-IN8 reverses the increase of proinflammatory activation of M2(IL-4) macrophages upon fucoidan treatment. Data representative of two independent experiments.

Figure EV7

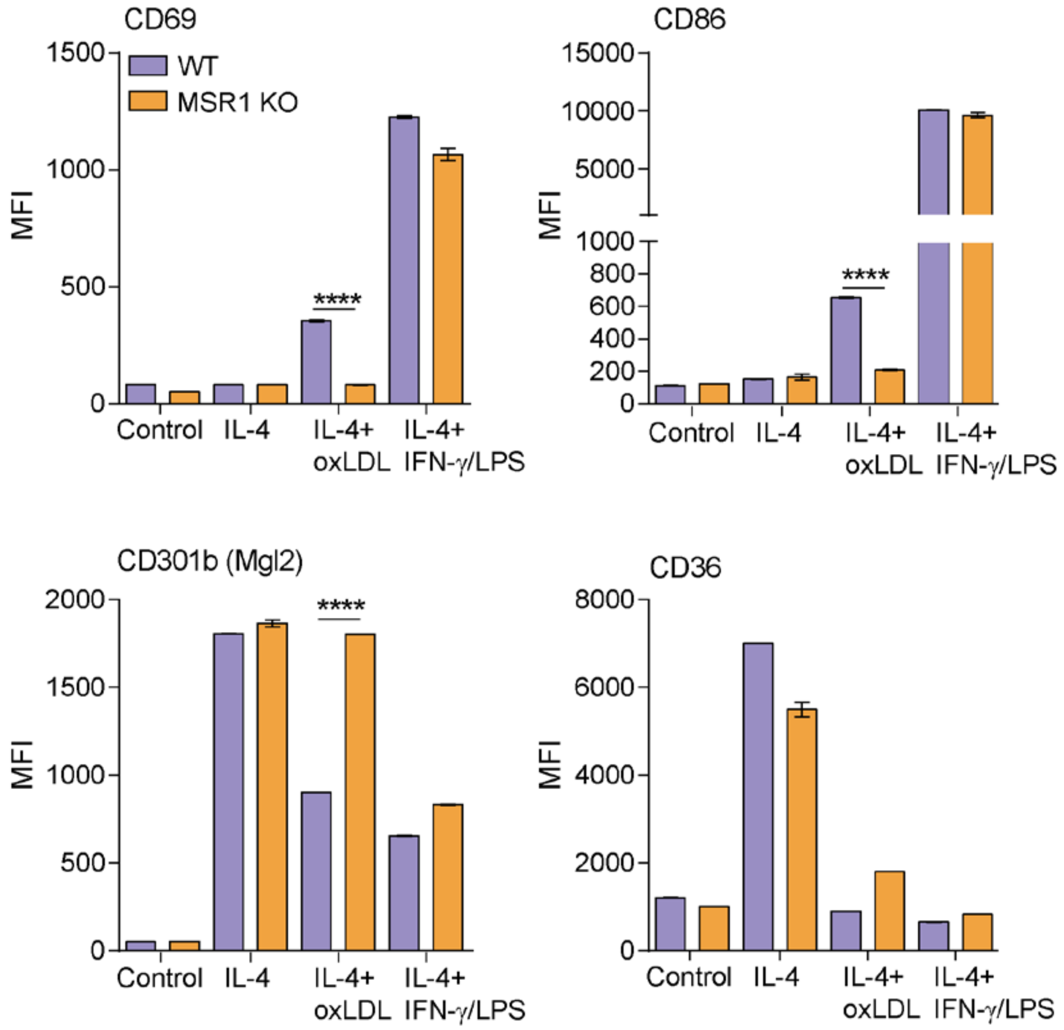


***Figure EV7: Quantitation of flow cytometry data in Figure EV6A.***

Data shows that triggering of MSR1 through oxLDL induces a more pro-inflammatory phenotype in IL-4 activated WT BMDMs but not in MSR1 KO macrophages. Data are shown as mean ± SEM from three independent experiments. MFI (Median Fluorescence Intensity).

### Figure EV8

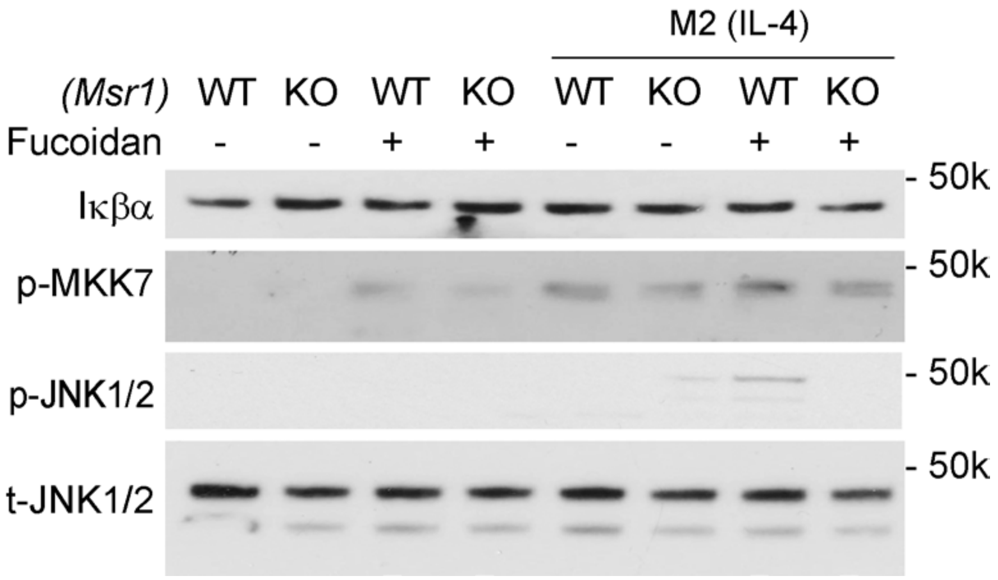


***Figure EV8: Triggering of MSR1 does not seem to activate NF-κB pathway.*** Immunoblot data shows that IκBα is not degraded in response to triggering of MSR1 with 50 µg/ml Fucoidan, indicating that the NF-κB pathway is not activated.
